# Absence of endosymbiotic gene transfer in symbiont-bearing diplonemids

**DOI:** 10.64898/2026.01.29.702485

**Authors:** Valéria Gabková Juricová, Vojtěch Žárský, Daria Tashyreva, Julius Lukeš, Aleš Horák

## Abstract

Horizontal gene transfer (HGT) is one of the fundamental processes in the evolution of prokaryotic genomes, while its importance in eukaryotes is still debated. Some of the hallmark eukaryotic organelles, such as mitochondria and chloroplasts, are of an ancient endosymbiotic origin. The process of acquiring (and losing) new endosymbionts is dynamic and still ongoing in many lineages. Endosymbiotic gene transfer (EGT) between symbionts and their hosts has been considered as one of the major sources of overall HGTs in eukaryotes. Thanks to recent advances in genomics and microscopy, more and more endosymbionts are discovered in protists offering suitable models for the study of EGT and its impact on the host. Recently, the presence of holosporacean and chlamydiacean symbionts in the novel strains of marine euglenozoan flagellates of genus *Rhynchopus* has been discovered. Here, we present an analysis of the genomes and transcriptomes of five *Rhynchopus* strains and examine the extent of EGT/HGT and the role of endosymbiosis in shaping the nuclear genome of symbiont-bearing hosts. Our results have shown that there is no evidence of a recent EGT from either Holosporales or Chlamydiales symbionts. The absence of such transfers suggests that EGT or at least a stable retention of EGT genes is not a requisite for a successful endosymbiosis. Furthermore, our results show striking differences between patterns of detected HGTs among the *Rhynchopus* lineages pointing to a dynamic and largely neutral evolution of horizontally-acquired genes.

## Introduction

Endosymbioses between microbial eukaryotes (protists) and prokaryotes in the marine environment are of growing interest for many studies (Tashyreva et al., 2018; Edgcomb et al., 2011; Balzano et al., 2015; Vannini et al., 2004; Maki et al., 2004). In protists, symbiotic interactions are either facultative or obligate and occupy a continuum between parasitism to mutualism (Husník et al. 2021). The exact nature of endosymbiotic relationships depends on both internal and external factors and can change on a temporal scale. In terms of negative effects, many symbionts are considered parasites or cause lytic infections (Thomas et al., 2006; Pagnier et al., 2015; Khalil et al., 2016). However, endosymbionts can also benefit the host by degrading toxins produced by prey organisms (Dirren et al., 2014), defend against bacterial and viral infections (Ishida et al., 2014;Arthofer et al.2022), synthesizing nutrients and vitamins (Leander and Keeling, 2004; Alves et al., 2013; Kostygov et al. 2017), or removing the excess of superfluous compounds (Gutiérrez et al., 2017; Hamann et al., 2016; Takeuchi et al., 2020). However, the role of the vast majority of symbionts in their host is still unknown (Husník et al. 2021).

A long-lasting, strictly host-associated lifestyle with selective pressure from the host system or strong genetic drift plays a role in the genome evolution of bacterial endosymbionts (McCutcheon and Moran, 2011; Latorre and Manzano-Marin, 2017). Their genomes usually undergo radical reduction in size and coding capacity, including an increase in pseudogenization and proliferation of selfish elements, and a decrease in the proportion of GC bases (Mira et al., 2001; Moran, 2002). Such a reduction often leads to the inevitable extinction of the symbiont or, in rare cases, when decreased protein-coding capacity is compensated by the import of host-encoded proteins, to its transformation into an organelle (Bennett and Moran, 2015; Keeling et al., 2015; Nowack 2014; Coale et al. 2024). Once the symbiosis is established, some of the endosymbiont’s genes are either lost or, far less commonly, incorporated into the host genome through endosymbiotic gene transfer (EGT) (Husnik and McCutcheon 2017; Henze et al., 2002). Although EGT has often been discussed as a key part of endosymbiotic integration, recent evidence suggests that this assumption is incorrect, and its extent, and general nature are debated (Hehenberger et al., 2016; Keeling and McCutcheon, 2017; Nikoh et al., 2010; Keeling, 2024).

Until recently, diplonemids, heterotrophic marine flagellates, were the only group within Euglenozoa without any known endosymbionts. This changed when the existence of alphaproteobacterial obligate symbionts in several diplonemid species was revealed (George et al., 2020; Prokopchuk et al., 2019; Tashyreva et al., 2018). Based on 16S rRNA phylogenies, *Diplonema japonicum* harbors symbionts from the order Holosporales, specifically *Cytomitobacter primus* and *Nesciobacter abundans*. *Diplonema aggregatum* is a host to *Cytomitobacter indipagum*. *Namystynia karyoxenos* from the Hemistasiidae contains *Sneabacter namystus*, a symbiont branching within Rickettsiales. Recently, phylogenomic pipeline implemented in GTDB database (https://gtdb.ecogenomic.org/) placed *Cytomitobacter* and *Nesciobacter*, together with holosporalean *Gromoviella agglomerans* and some rickettsialean metagenomes as an independent lineage within Rickettsiales. However, this branching lacks support and further evidence justifying change of diplonemid symbionts classification is needed. Holosporales and Rickettsiales are characterized by highly streamlined genomes and their metabolic capacities are often extremely reduced (Castelli et al., 2022; Garushyants et al., 2018; Floriano et al., 2018). Greatly reduced genomes were also observed in diplonemid symbionts. This, along with comparable gene content and overall predicted metabolic potential, indicates convergent evolution of the alphaproteobacterial endosymbionts of diplonemids (George et al., 2020). They are located in the host cytoplasm or the nucleus, and some of them closely interact with the mitochondrion (Prokopchuk et al., 2019; Tashyreva et al., 2018). In addition to the previously mentioned occurrences, the holosporacean symbionts were also recently discovered in two novel strains of *Natarhynchopus humris*, whereas the two novel strains of *Rhynchopus euleeides* contain *Syngnamydia salmonis* (Simkaniaceae; Chlamydiales) endosymbiont. Based on 16S rRNA analysis, this chlamydial symbiont of diplonemids is closely related to major fish pathogens that cause epitheliocystis and to endosymbionts of amoebae. (Nylund et al., 2018; Fehr et al., 2013; Tashyreva et al. 2025). All these discoveries indicate that diplonemids have a great diversity of endosymbionts. Although holosporacean symbiont phylogenies are broadly congruent with those of their diplonemid hosts (Tashyreva et al., 2018, 2025), suggesting their coevolution, the patterns of symbiont presence and absence allow for alternative hypotheses such as independent gains or horizontal transfers of the symbionts among the co-occurring diplonemid species (Tashyreva et al., 2018). Their role and influence on host metabolism also remain unclear.

In prokaryotes, where the HGT is one of the major processes in genome evolution, the best studied and widely occurring mechanisms are conjugation, transduction via bacteriophages, and the movement of mobile genetic elements such as transposons (Filée, 2014; Schaack et al., 2010; Koraimann and Wagner, 2014). Very important and widely discussed is for example the antibiotic resistance of bacteria acquired via HGT, leading to lethal strains such as vancomycin and methicillin-resistant *Staphylococcus aureus* (Chang et al., 2003).

While in eukaryotes the importance of HGT is still debated, their ecology provides additional opportunities like the HGT from phagocytosed cells and from the endosymbionts via EGT (Keeling and Palmer, 2008; Husnik and McCutcheon, 2018; Van Etten and Bhattacharya, 2020). In many cases, HGT plays an important role in adaptation of eukaryotes to different environmental conditions. As an analogy to the antibiotic resistance of prokaryotes, HGT is an important mechanism shaping the evolution of parasitic protists such as *Toxoplasma*, *Blastocystis* or *Entamoeba* (Romero et al., 2016; Angel et al. 2024; Eme et al., 2017), giving them the ability to adapt to the host and evade its immune responses.

Here we analyze genome assemblies of three *Rhynchopus* and two *Natarhynchopus* strains, supplemented by transcriptome sequencing to investigate: 1) the role of endosymbiosis in the context of shaping the nuclear genome of the symbiont-bearing hosts by EGT, and 2) the acquisition of horizontally transferred genes from donors other than the current symbionts and the extent of metabolic capabilities they confer.

## Materials and Methods

### Cell Cultures

*Rhynchopus asiaticus*, *Natarhynchopus humris* KQ11, *Rhynchopus euleeides* FC903 and YPF1915 strains were isolated from seawater samples collected in coastal waters of the Sea of Japan. Strain *N. humris* DT0507 was isolated from a water sample from Enoshima aquarium (Kanagawa, Japan). Clonal axenic cultures were established by manual picking of single cells with a glass microcapillary and inoculating into Hemi medium (Tashyreva et al. 2018), supplemented with 10 µl/ml antibiotics cocktail (P4083, Sigma-Aldrich). All strains were subsequently cultivated axenically in Hemi medium at 13°C without the addition of antibiotics.

### DNA isolation and sequencing

High molecular weight total DNA was isolated by phenol-chloroform extraction as follows. Pelleted cells (50 mg) were resuspended in 1.5 ml lysis buffer (100 mM Tris-HCl pH 8.0, 100 mM EDTA pH 8.0, 1.15 % polyvinylpyrrolidone, 5 mM spermidine, 0.5% N-lauryl sarcosine), treated with 0.2 mg of RNAse for 30 min at 37°C and with 100 μL proteinase K (10 mg/mL) for 60 min at 37°C, mixed with an equal volume of Phenol: Chloroform: Isoamylalcohol (25:24:1) and centrifuged at 4,000 g and 4°C for 10 min. Aqueous phase was mixed with 2 volumes of ice-cold ethanolic solution of 0.5 M ammonium acetate. Precipitated DNA was removed with a glass hook, rinsed with 1 mL of pre-chilled 80% ethanol (4°C) and solubilized in 50 μL of 10 mM Tris-HCl pH8.5 at 4°C overnight. A TruSeq DNA libraries were analyzed using Illumina Hiseq 2500 to generate 99.56, 105.27, 96.03, 95.57, and 90.05 million 150 bp paired-end reads.

### RNA isolation and sequencing

For transcriptomics, cultures were grown in triplicates in horizontal plastic flasks and harvested in the late logarithmic phase of growth (∼9·10^5^ cells/ml for *R. asiaticus*). Total RNA was isolated and purified from DNA with Master Pure Complete DNA and RNA purification kit (Lucigen, Cat. N. MC85200) following the manufacturer’s protocol.

Poly(A) enriched RNA libraries were sequenced using Illumina Hiseq 2500 to generate 150bp paired-end reads for *R. asiaticus* (31.02, 26.24, 31.22 mil.), *N. humris* DT0507 (57.46, 42.26, 48.36 mil.), and *N. humris* KQ11 (62.03, 53.93, 48.43 mil.).

### Transcriptome assembly and quantification of transcripts expression

Evaluation of the quality of data was performed using FastQC v0.11.9 (Andrews, 2010). Filtering of low-quality reads and adapters was done using Trimmomatic v0.39 (Bolger et al. 2014). Transcriptome assembly was obtained using rnaSPAdes v3.15.1 (Bushmanova et al. 2019) with the default parameters. Protein sequences were identified with TransDecoder v5.5.0 (Haas), HMMER (Eddy, 2011) and DIAMOND v2.0.7 (Buchfink et al. 2015). The expression of transcripts was quantified and normalized using Salmon (Patro et al., 2017).

### Genome assembly and annotation

The low-quality short reads and adapter sequences were trimmed using Trimmomatic v0.39 (Bolger et al. 2014) and fastp (Chen et al. 2018). The read quality before and after trimming was monitored with FastQC (Andrews, 2010). Cleaned read sets were assembled in SPAdes v3.15.1 (Bankevich et al. 2012). Contigs shorter than 300 bps were discarded. For read representation estimation, bwa v0.7.017 (Li et al. 2009) and samtools v1.10 (Danecek et al. 2021) were used to map reads back to assemblies. To assay the composition of the sequenced cultures, SSU rRNA sequences were detected using Infernal (Nawrocki and Eddy, 2013) and compared with the SILVA rRNA database and NCBI nt database. To remove contamination and classify assembled sequences (host or symbiont), contigs were searched against the NCBI nt and UniProt Reference Proteomes database using the BLASTN and DIAMOND protein aligner v2.0.7 in sensitive mode (Buchfink et al. 2015). The contigs were classified based on blast matches to multiple loci in a semi-supervised manner using in-house script (https://github.com/vojtech-zarsky/vojta-tools/blob/master/classifyContigs.py). The quality of the host genomes was assessed using Quast (Gurevich et al., 2013). In strains with both genomic and transcriptomic data available, the transcriptomic reads were mapped on the host contigs using splice-aware aligner TopHat (Trapnell et al. 2009). Gene prediction was performed using GeneMark-EX and Augustus within BRAKER (Bruna et al. 2021; Hoff et al. 2019). The quality and completeness of the genome assembly and gene prediction were examined using Benchmarking Universal Single-Copy Orthologs (BUSCO) v. 5.7.1 using eukaryota_odb 10 lineage dataset (Manni et al. 2021). Gene annotation was performed using InterProScan (Jones et al. 2014), OmicsBox (Conesa et al. 2004) and eggNOG-mapper (Cantalapiedra et al. 2021). Gene ontology analysis and Clusters of orthologous groups (COG) annotation were performed to infer the biological processes associated with the identified HGTs.

### Molecular taxonomy

To place so-far undescribed diplonemid isolates used in this study in the diplonemid taxonomic context, we have extracted 18S rRNA sequences from genomic assemblies (see above) and included them in the dataset used in Tashyreva et al. (2025), aligned using MAFFT v7.505 (Katoh & Standley, 2013). Ambiguous and hypervariable sites were then removed in SeaView 5 (Gouy et al., 2021) resulting in alignment containing 85 species and 1793 positions. Phylogenetic placement of new diplonemids was inferred using the maximum likelihood under the SYM model with four free-rate categories (SYM+R4) as implemented in IQTree 2 (Minh et al., 2020). This particular model was found to be the best-fitting according to both Akaike and Bayesian information criterion as inferred by model finder implemented in IQTree 2. The branching support was estimated using the non-parametric bootstrap values inferred from 500 replicates in IQTree 2 (-b 500 option).

To build a protein supermatrix for phylogenomic inference, orthologs of 20 conserved genes were collected from the predicted diplonemid proteomes and from other selected euglenozoans and *Naegleria gruberi* (Table S1). Each gene set was aligned using MAFFT and the alignment was trimmed using BMGE. The trimmed alignments were then merged into a species supermatrix containing 10 species and 6866 positions. A maximum likelihood phylogenetic tree was inferred using the LG+PMSF+G model (Wang et al., 2018) implemented in IQTree 2. The branching support was estimated using 10000 replicates of ultrafast bootstrap (Hoang et al., 2018).

### Phylogenomic analysis

For each of the diplonemid protein coding gene phylogenies were estimated. In the first step, the protein sequences were searched against the UniProt and EukProt databases using DIAMOND (Buchfink et al. 2015) in sensitive mode. The protein sets of all diplonemids were also compared with each other to detect homologs within these species. All hits with e-values higher than 1e-5 were discarded. To minimize redundancy, highly similar sequences were reduced using cd-hit with the 90% identity cutoff. A multiple sequence alignment was constructed for each dataset using MAFFT (Katoh and Standley, 2013) with the default settings and trimmed with BMGE (Criscuolo and Gribaldo, 2010) using the BLOSUM30 substitution matrix. Phylogenetic trees were inferred with FastTree (Price et al. 2009) with support values estimated from the minimum of SH-like and approximate likelihood ratio test supports.

## Results and Discussion

### Genomes

Five genomes were sequenced using Illumina HiSeq 2500 platform and assembled using SPAdes. The three *Rhynchopus* assemblies range between 105-113 Mb and are composed of many contigs (∼29-39k) which is reflected by low N50 values between 4.8-7.1 kbs. The two *N. humris* assemblies appear to be more compact with sizes between 60-64 Mb lower number of contigs (∼7-12k) and better N50 values between 16.1-17.8 kbs (Fig. 1B). A rough estimate of genome size for *Rhynchopus* species ranges from 200-300 Mb.

**Fig. 1.**
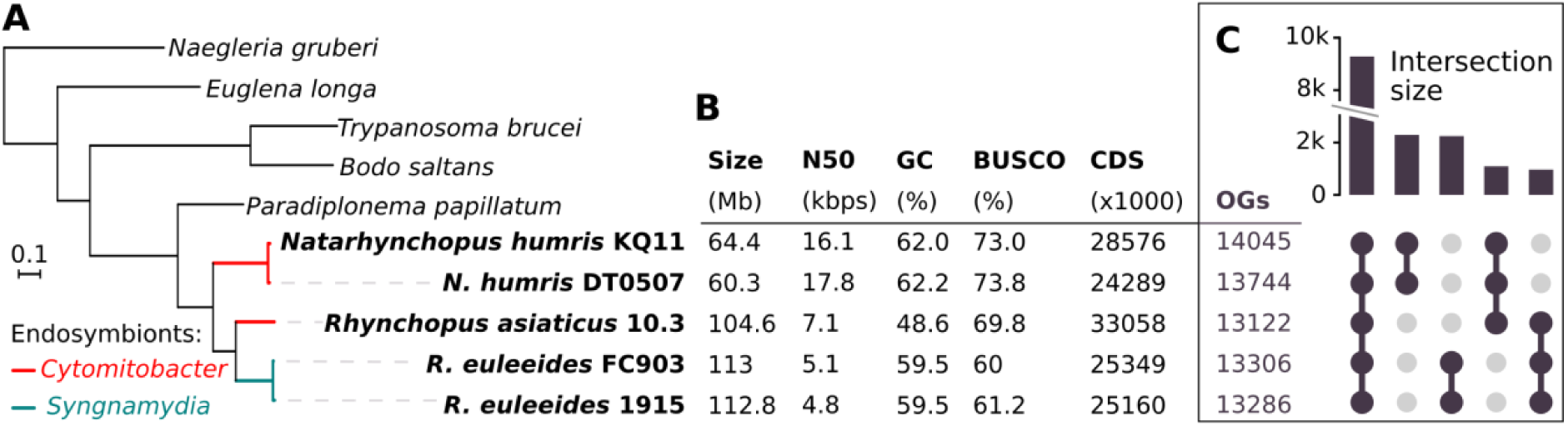
(A) Maximum-likelihood phylogeny of the diplonemid representatives based on 20 conserved nuclear-encoded genes (10 taxa, 6866 aligned positions) under the LG+PMSF+G4 model as inferred by IQTree2. All the nodes received a full support from 10000 ultra-fast bootstrap replicates. Lineages hosting *Cytomitobacter* and *Syngnamydia* endosymbionts are shown in red and green respectively. (B) Parameters of assemblies of diplonemid genomes used in this study. (C) An UpSet plot representing the sum of all orthologous groups (OGs) per lineage and shared orthologous groups of the diplonemid representatives as predicted by OrthoFinder. Only the five most common intersections (patterns of shared OG) are shown.

The SSU rRNA analysis and the contigs classification confirmed the identity of the hosts and their endosymbionts, and showed a very low level of contamination. Each diplonemid host possesses one bacterial symbiont, with the exception of *N. humris* KQ11. This strain is a host to two *Cytomitobacter* endosymbionts, one of which is also present in *N. humris* DT0507.

The quality and completeness of the genome assemblies were examined using the BUSCO assessment (Simão et al. 2015). Completeness scores ranged from 60,0 % to 73,8 %. We perceive these values to be good, considering that there are no closely related representatives in the BUSCO database. For comparison, using the same methodology, the predicted proteome of the relative *Paradiplonema papillatum*, the only diplonemid with annotated genome published (Valach et al., 2023) has BUSCO score of 68.6%.

The protein-coding genes were predicted with the aid of transcriptomic data in three strains (*N. humris* KQ11 and DT0507, and *R. asiaticus*). In the two strains without transcriptomic data (*R. euleeides* 1915 and FC903), proteins from the previous three predictions were used as a guide for gene prediction using BRAKER (Bruna et al. 2021). The missing transcriptomes for *R. euleeides* could have contributed to their lower BUSCO scores (60.0, 61.2%) due to reduced annotation accuracy.

### The HGT analysis revealed no evidence for EGT and a high turnover of transferred genes

We developed a two-step pipeline of HGT analysis to detect and characterize EGT/HGT in host genomes: 1) a comprehensive semi-automatic phylogenetic analysis of all genes in five diplonemid genomes was performed, together with a thorough manual inspection of each phylogenetic tree in a refined set to identify a HGT signal and confirm/refute a possibility of EGT; 2) annotation of all genes using InterProScan, OmicsBox and eggNOG-mapper to determine their function.

We have found 93 candidates for horizontally transferred genes of prokaryotic origin in the nuclear diplonemid genomes. Some of them contain spliceosomal introns all of which have canonical splice-site boundaries (GT/AG), however, on average, the HGT candidate genes had much lower number of introns compared to the rest of nuclear genes (2.7-7.6-times less, Table_S4.Intron_counts.xlsx). The topologies of the curated set of putative HGT genes were thoroughly analysed to infer putative prokaryotic donor taxa and monophyly with some other eukaryotic groups. Strikingly, there was a very small overlap of HGTs shared between the three diplonemid species (Fig. 2A), pointing to a high evolutionary turnover and predominantly neutral evolution of HGTs in this system. More than half of the putative diplonemid HGTs of the were shared with other eukaryotic lineages outside of Euglenozoa (51). We found the predominantly marine ochrophytes, dinoflagellates and haptophytes to be most commonly occurring as orthologous with the diplonemid HGT candidates (Fig. 2B). This points to probable transfers between co-occurring eukaryotic lineages.

**Fig 2.**
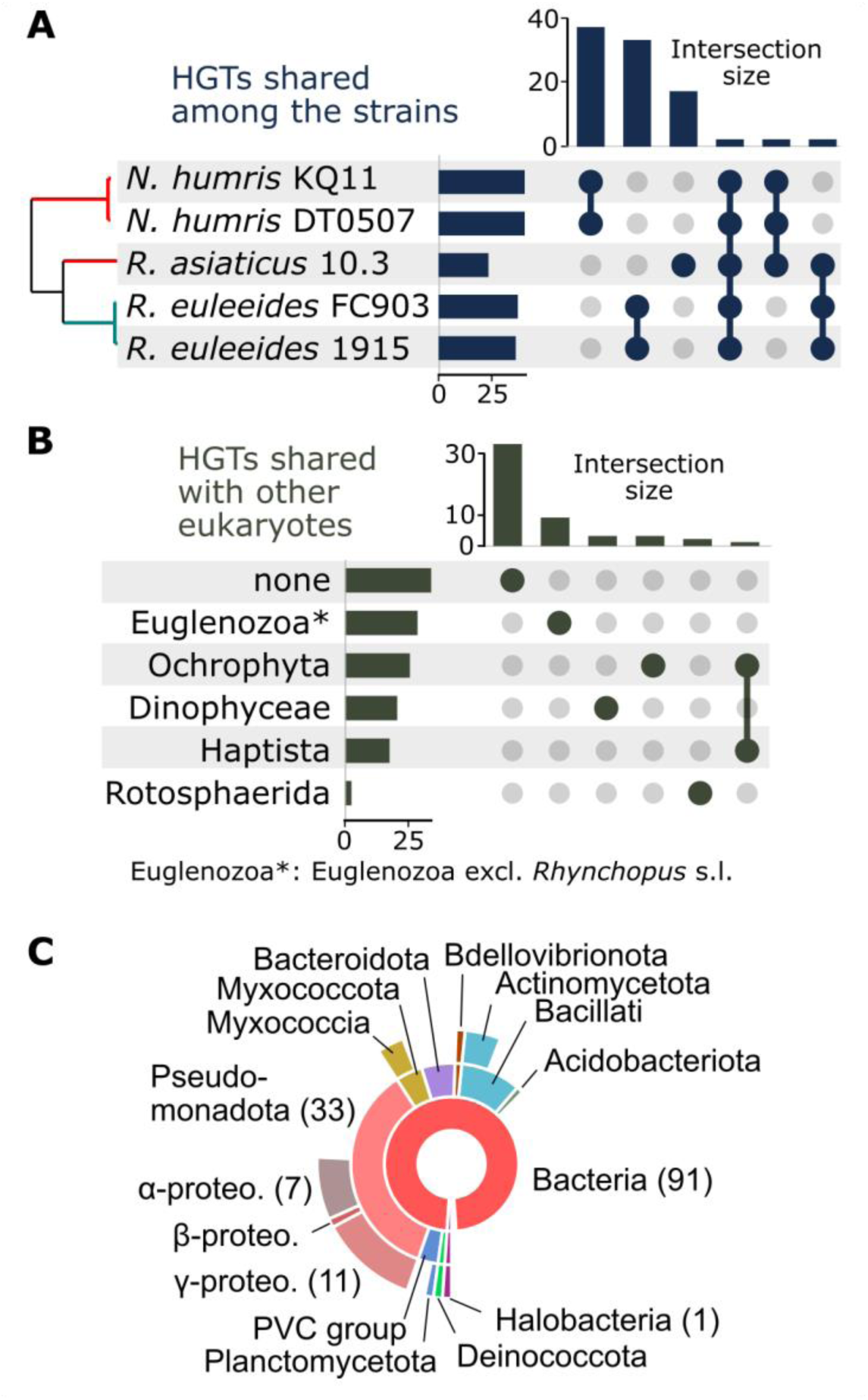
Analysis of 93 putative horizontal gene transfers (HGTs). (A) An UpSet plot of the HGTs shared between the five diplonemid strains. (B) An UpSet plot representing selected eukaryotic lineages forming a monophylum with the putative HGTs and the six most common combinations (intersections). The group Euglenozoa* represents Euglenozoa with the exclusion of the *Rhynchopus* and *Natarhynchopus* lineages (C) A pie chart representing inferred sources of the putative prokaryota-eukaryota HGTs. The chart is nested to represent the taxonomical hierarchy of given prokaryotic groups. For one gene, the source was not inferred, for another gene an archaeal lineage was identified and for the remaining 91 HGTs bacteria were identified as the source.

We could not find any evidence of diplonemid host sequences branching within a particular endosymbiotic clade (Holosporales, Chlamydiales), nor of symbiont sequences integrating into the host genome; therefore, no recent EGT signal was detected. However, some of the putative HGTs branched within Pseudomonadota (formerly Proteobacteria) (33), mostly free-living Gamma- and Alphaproteobacteria, while for many (34) we weren’t able to pinpoint an origin within certain taxon (Fig. 2C). It is therefore possible that the analysed hosts, and probably other diplonemids may have harboured transient proteobacterial endosymbiont(s) in the past, however other modes of HGT, e.g. from food, are certainly possible. These findings lead to the proposal that the acquisition of genes by the nucleus happened in accordance to the „shopping bag” model (Larkum et al., 2007).

Although a growing number of Holosporales genomes have become available, our understanding of their genomics and evolution remains limited. According to available genomic data, many of these so-called “professional symbionts” often exhibit varying degrees of genome reduction, sometimes as extreme as approximately 600 kb genome, and live in association with protists (George et al., 2020; Castelli et al., 2022; Shiohama et al., 2022). Their intracellular lifestyle likely originated in their early evolutionary history, followed by the horizontal transfer of symbiont species between hosts over time (Wang and Luo 2021; Hugoson et al. 2022). Even though these symbionts are host-dependent, they are not strictly confined to one host species and exhibit a wide range of host specificity, from narrow specialists to broad generalists. The question remains as to how reduced the genome of the common ancestor of diplonemid symbionts was and what degree of reduction *Cytomitobacter* or other Holosporales have experienced in diplonemid hosts. The absence of EGT signal suggests that even if some genome downsizing did occur in diplonemids, other factors must have been a driving force leading to it. Whether genetic drift or trophic specialization played a role or the genes were simply lost remains unclear (Lee and Marx, 2012; Qin et al. 2019).

The absence of EGT from *Chlamydiales* endosymbionts in host genomes is probably much easier to explain, as genome evolution in some *Chlamydiales* also includes gene acquisition, rather than being limited to gene loss alone (Dharamshi et al., 2023). Members of the phylum Chlamydiae are ubiquitous in the environment.

Some families of Chlamydiales are known to have a specific natural host (e.g., Waddliaceae, Parachlamydiaceae), but most species exhibit an extremely broad host range, with frequent host species jumps (Collingro et al., 2011; Read et al., 2013). This lifestyle requires evolutionary adaptations that allow the bacteria to survive outside host cells during extracellular transmission and to successfully infect new host species.

Therefore, gene transfer should be kept to a minimum to avoid genome reduction of the bacterial symbiont/parasite. This could be the reason why we could not find any traces of the putative EGT signal in the host genome.

Additionally, these Chlamydiales symbionts encode a substantially broader metabolic capacity than Holosporales, suggesting that they may retain a greater degree of functional independence and exhibit reduced reliance on host-derived genes.

Another hypothesis could be that *Syngnamydia* acted not as a donor but as a recipient of genes in this particular symbiosis/host relationship, however except for more general eukaryotic-like genes in the *Syngnamydia* genome we didn’t find any support for such recent transfer.

### Function of HGTs

Although HGT analysis did not confirm an EGT signal in any of the analyzed host genomes examined, we were able to detect the presence of at least 93 HGTs of bacterial origin, 84 of which could be functionally annotated (Table S2). The breakdown of the functions of HGT genes in the COG functional categories showed an enrichment of functions related to carbohydrate metabolism and transport (22), followed by genes involved in amino acid metabolism (10), energy metabolism (9), and lipid metabolism (5) (Fig. S2). Our results are consistent with previous findings that genes of carbohydrate metabolism and energy conversion are frequently transferred between free-living microorganisms (Song et al., 2019). We have identified four HGT-CAZyme families, comprising 13 members in total. Specifically, the most members were identified in the glycoside hydrolase family (5), followed by the carbohydrate-binding module family (3), the polysaccharide lyase family (3), and finally the carbohydrate esterase family (2). One of the interesting genes that have emerged as HGT from the bacterial phylum, particularly the Pseudomonadota group, is hydroxymethylglutaryl-CoA synthase (HMGCS) (2.3.3.10). Notably, this gene was only identified in strains DT0507 and KQ12, suggesting selective acquisition or retention rather than a universally shared HGT within the *Rhynchopus/Natarhynchopus* group. This synthase is the key enzyme of the mevalonate pathway, which produces various precursors for isoprenoid compounds (Edwards and Ericsson, 1999). The ability to produce isoprenoid precursors could give the host a metabolic advantage by enabling the synthesis of essential compounds such as sterols (e.g. cholesterol), isoprenoids and dolichols, which are important for cell signaling, membrane structure and other cellular processes. Isoprenoids play a role in stress responses, suggesting that the diplonemid host may utilize this metabolic pathway to adapt to environmental challenges.

Another intriguing horizontally transferred gene from Pseudomonadota group is thebaine 6-O-demethylase. This enzyme is primarily associated with plants, especially in the biosynthesis of opiate alkaloids, but the gene has also been found in genomes of several bacteria, and the dinoflagellate Symbiodinium (Aranda et al., 2016; Zhan and French, 2019). Through its action on thebaine, thebaine-6-O-demethylase contributes to the production of important opiate precursors such as oripavine. This is a crucial step in the formation of morphine and codeine, which are important medicinal compounds. A closer examination of the genomes of *Rhynchopus* did not reveal presence of any other genes involved in the biosynthesis of phytochemical compounds, so the role of this enzyme in diplonemids remains a mystery. In the host genome, we have also observed the presence of HGT involving a gene linked to antibiotic metabolism. Penicillin amidase (3.5.1.11), a member of the penicillin acylase family, is associated with the hydrolysis of penicillin, a widely used antibiotic. This enzyme is mainly found in bacteria and fungi, but the gene has also been found in the genomes of several protists (Clarke et al., 2013; Liechti et al., 2019). Both thebaine 6-O-demethylase and penicillin amidase, as well as many other genes were detected only in *R. euleeides* strains, again pointing to genus-specific HGT.

We also found the gene encoding putrescine aminotransferase (PATase) (2.6.1.82), which catalyzes the putrescine degradation reaction. In eukaryotes, putrescine is utilized by spermidine synthase to produce spermidine, the intermediate of spermine. In bacteria, putrescine can be converted by PATase to 2-oxogutarate (Prieto-Santons et al., 1986) and finally to 4-aminobutanoate, allowing bacteria to utilize putrescine as their sole source of nitrogen. This enzyme and metabolic pathway are virtually absent in eukaryotes. Our phylogenetic analysis suggests an acquisition from bacteria to the *R. euleeides* strains, as this protein clusters within a group of bacterial homologs. Since the oxidative deamination of putrescine, and subsequent oxidation of GABA aldehyde leads to the production of 4-aminobutanoate (GABA), it can be speculated that the presence of PATase would be advantageous for the diplonemid host for the alternative pathway of GABA production. Considering that GABA functions both as an intracellular signal and a messenger in protozoan parasites, it is plausible that horizontally acquired GABA-related genes in the host genome could play a role in mediating or modulating diplonemid interactions with their multicellular hosts (Nugraha et al., 2022).

A surprising discovery was that of a homologue of a bacterial haemolytic enterotoxin in the *N. humris* genomes. Horizontal gene transfer of bacterial toxin genes to eukaryotes is a rare event, but several cases have been described (Verster et al., 2021; Moran et al., 2012). The acquisition and utilization of bacterial toxin genes by eukaryotes can provide considerable functional benefits, especially in defense mechanisms and interactions with other organisms (Chou et al., 2014; Ambrose et al., 2014; Moran et al., 2012). A diplonemid, very closely related to *N. humris* DT0507 and KQ11 (>99% 18S rRNA identity), was previously isolated directly from the blood of lobsters (von der Haiden 2004; Tashyreva et al 2022). The toxin may therefore support the parasitic lifestyle of *N. humris* by promoting the invasion and persistence of diplonemids within host blood and tissues by disrupting their cellular integrity. Parasitic and predatory behavior has been commonly observed in diplonemids (Tashyreva et al. 2022), therefore, the gene could facilitate interaction with the host when diplonemids engage in parasitic relationships, which could allow them to gain nutrients or evade the immune system.

Our results are broadly comparable to those reported in Valach et al. (2023),), with partial overlap in the functionality of HGT. Similarly to their findings in *Paradiplonema papillatum*, HGTs in the analysed diplonemids were dominated by metabolic enzymes, including oxidoreductases, hydrolases, or lyases, and are enriched in CAZymes. We have also identified genes involved in redox balance and detoxification, such as glutathione S-transferase.

### Several metabolic pathways are composed of genes of mosaic origin

A function was assigned to 73 of 93 HGTs. The majority of genes were identified as metabolic enzymes. Hydrolases (29) which are responsible for degradation reactions in the cell, were most frequently represented. In second place were oxidoreductases (18), transferases (10) and lyases (9). As far as the acquisition of HGT genes is concerned, the results do not show a consistent pattern in relation to the source. In most cases, a metabolic pathway had several different donor taxa.

This results in is an observed mosaic evolutionary history of metabolic pathways, some of which were affected by several cases of HGT. Out of 93 HGTs, the most enriched pathway of carbohydrate metabolism, with 7 HGTs found in *R. asiaticus*, 9 in *R. euleeides* strains, and 3 in *N. humris* strains. Four enzymes were involved in central carbon metabolism, in particular glycolysis, tricarboxylic acid cycle (TCA), and the pentose phosphate pathway. In case of these specific pathways, the acquisition of four HGTs from Pseudomonadota and the FCB group in *R. euleeides* probably conferred the ability to metabolize cyclodextrins. These genes encode enzymes that sequentially degrade cyclodextrin to maltodextrin, maltose and finally D-glucose, enabling the host to utilize complex carbohydrates from starch and glycogen (Fig. 3). The transferred genes correspond to important hydrolases such as cyclodextrinase (EC 3.2.1.54) and α-glucosidases (EC 3.2.1.20), which are crucial for the degradation of oligosaccharides. This metabolic capacity would help *R. euleeides* to utilize polysaccharides and oligosaccharides from the environment more efficiently, which could provide a competitive advantage in carbohydrate-rich niches. Besides opportunistic parasitism, diplonemids are known for predation on large planktonic microorganisms, such as diatoms and dinoflagellates, and small crustaceans (copepods), as well as invasion of freshwater and marine plants (Tashyreva et al. 2022; Valach et al. 2023). These prey organisms contain diverse polysaccharides as components of their cell walls and exoskeletons, as well as storage products.

**Fig 3.**
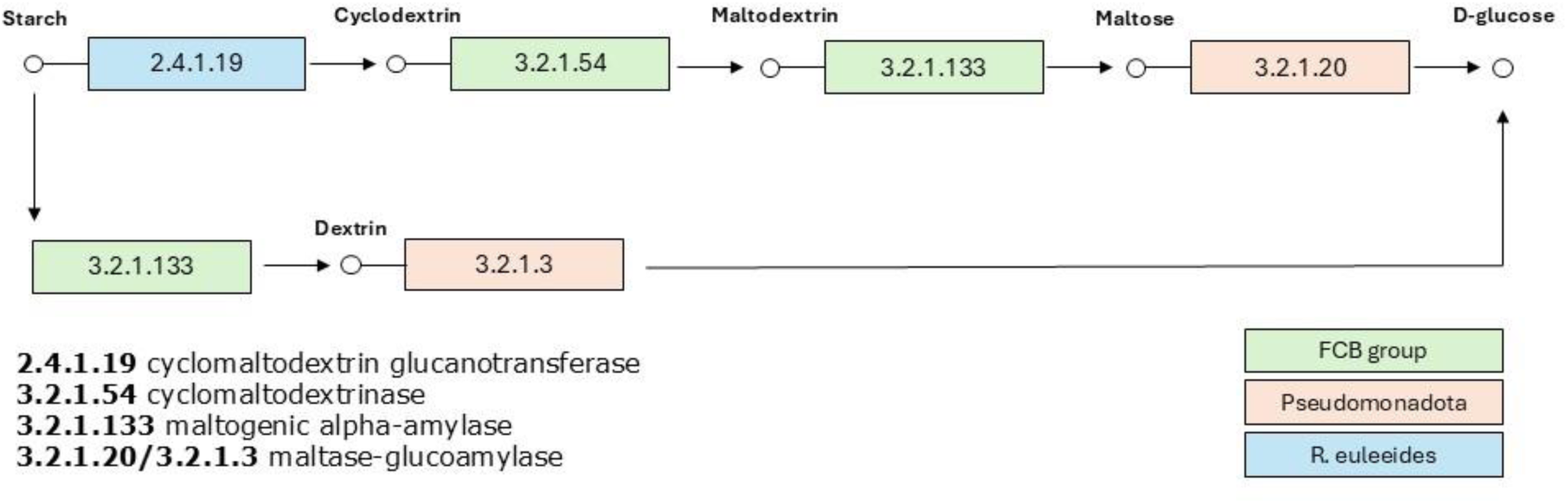
Starch degradation in *R. euleeides* is enabled by horizontally acquired genes from Pseudomonadota and the FCB group. These genes encode key enzymes such as cyclodextrinase and α-glucosidase, which sequentially convert cyclodextrins into maltodextrins, maltose and finally D-glucose. Cyclomaltodextrin glucanotransferase is of eukaryotic origin.

14 HGTs have been assigned to amino acid metabolic pathways, most of which (5) are involved in glycine, serine and threonine metabolism. The acquired genes include transferases (aminomethyltransferase, glycine C-acetyltransferase), lyases (cystathionine gamma-lyase, L-threonine dehydratase) and sarcosine oxidase. More specifically, 8 HGTs involved in amino acid metabolism are scattered in the *N. humris* strains — typically occurring in the form of one or two genes per metabolic pathway. This fragmented distribution suggests that *N. humris* has not acquired entire metabolic pathways through HGTs, but rather isolated genes that may complement or enhance existing functions. This pattern — single, seemingly random HGT events rather than a coordinated acquisition of entire metabolic pathways — is also observed in other metabolic pathways beyond amino acid metabolism. It suggests a more opportunistic or piecemeal evolutionary strategy in which *N. humris* integrates useful genes as they become available. The metabolic pathways most affected in *R. asiaticus* were carbohydrate metabolism, particularly fructose and mannose metabolism, amino acid metabolism, methane metabolism, and nicotinate and nicotinamide metabolism. The distribution of HGTs in the metabolic pathways was mosaic-like, as in the other hosts. One of the interesting species-specific horizontal transfers of Pseudomonadota origin (to only *R. euleeides* strains) is gene coding 4-carboxymuconolactone decarboxylase in bacteria. This enzyme is involved in degradation of aromatic compounds, specifically benzoate. Analysis of the enzymes acquired by horizontal gene transfer revealed that most of them are involved in the catabolic rather than the anabolic parts of the metabolic network.

Transcript quantification from RNA-seq data of three strains (*R. asiaticus* 10.3, *N. humris* DT0507 and KQ11) revealed that the most highly expressed genes include various ribosomal proteins and histones. We detected the expression of all HGTs in the transcriptomic datasets. After normalisation, the expression profiles of the HGTs did not differ significantly from those of the native genes, indicating similar transcriptional activity. However, the expression levels among the HGTs varied, especially genes such as alpha-glucosidase, alpha-galactosidase or L-threonine dehydratase were highly expressed (Table S3).

## Conclusion

Our comprehensive genome and phylogenomics analysis of five diplonemid strains provides valuable insights into the ongoing dynamics of endosymbiosis and horizontal gene transfer in diplonemids. Although we found no clear evidence of an endosymbiotic gene transfer from the Holosporales or Chlamydiales symbionts, our study describes the complex role of prokaryotic horizontal gene transfer in shaping host organisms’ genetic makeup. The presence of 93 horizontally transferred prokaryotic genes—often shared with other marine eukaryotic lineages—suggests that environmental factors and interactions with surrounding microbial communities contribute to the evolution of marine protists. Strikingly we found very little shared HGTs among the three diplonemid species irrespective of their phylogenies, pointing to a very dynamic and rather neutral pattern of HGT gain and loss. HGTs from different bacterial taxa enhance or alter host metabolic capacities and primarily improve catabolic metabolic pathways. These results emphasize the importance of understanding the mechanisms and the broader ecological context in which EGT/HGT occurs and the evolutionary implications for both symbionts and their eukaryotic hosts. Diplonemids are a large emerging group of ecologically important marine protists, and the recent research only begins to unveil the vast diversity of this group and of their endosymbionts. Certainly, much more is to be learnt from them.

## Acknowledgement

We thank Patrick J Keeling for valuable insights and discussions. We acknowledge the support from the Czech Science Foundation (21-26209S) to V. G. J., D. T. and A. H. and the Gordon and Betty Moore Foundation (GBMF #9354) to V. Ž., and J.L. V. Ž. was further supported by the European Union under the LERCO project number CZ.10.03.01/00/22_003/0000003 via the Operational Programme Just Transition.

## Conflict of Interest statement

The authors have no conflict of interest.

## Data availability

All datasets supporting the conclusions of this article are included within the article or as additional files. The raw DNA-Seq, and RNA-Seq reads have been deposited in the European Nucleotide Archive (ENA) under accession number PRJEB106601 including genome and transcriptome assemblies.

## Author contributions

VGJ, VŽ and AH designed research; VGJ and VŽ performed research and analyzed data; DT cultivated cells; and VGJ, VŽ, DT, JL and AH wrote the manuscript.

## Notes

### Competing Interest Statement

The authors have declared no competing interest.

## References

Alves JM, Serrano MG, Maia da Silva F, Voegtly LJ, Matveyev AV, Teixeira MMG, Camargo EP, Buck GA. 2013. Genome evolution and phylogenomic analysis of Candidatus Kinetoplastibacterium, the betaproteobacterial endosymbionts of Strigomonas and Angomonas. Genome Biol Evol 5:338–350.

Ambrose KV, Koppenhöfer AM, Belanger FC. 2014. Horizontal gene transfer of a bacterial insect toxin gene into the Epichloë fungal symbionts of grasses. Sci Rep 4:5562.

Andrews S. 2010. FastQC: A Quality Control Tool for High Throughput Sequence Data. http://www.bioinformatics.babraham.ac.uk/projects/fastqc/

Angel SO, Vanagas L, Alonso AM. 2024. Mechanisms of adaptation and evolution in Toxoplasma gondii. Mol Biochem Parasitol 258:111615.

Aranda M, Li Y, Liew Y, et al. 2016. Genomes of coral dinoflagellate symbionts highlight evolutionary adaptations conducive to a symbiotic lifestyle. Sci Rep 6:39734.

Arthofer P, Delafont V, Willemsen A, Panhölzl F, Horn M. 2022. Defensive symbiosis against giant viruses in amoebae. Proc Natl Acad Sci USA 119(36):e2205856119. 10.1073/pnas.2205856119

Balzano S, Corré E, Decelle J, et al. 2015. Transcriptome analyses to investigate symbiotic relationships between marine protists. Front Microbiol 6:98.

Bankevich A, Nurk S, Antipov D, et al. 2012. SPAdes: a new genome assembly algorithm and its applications to single cell sequencing. J Comput Biol 19(5):455–477. 10.1089/cmb.2012.0021

Bennett GM, Moran NA. 2015. Heritable symbiosis: the advantages and perils of an evolutionary rabbit hole. Proc Natl Acad Sci USA 112:10169–10176.

Bolger AM, Lohse M, Usadel B. 2014. Trimmomatic: A flexible trimmer for Illumina sequence data. Bioinformatics 30:2114–2120.

Bruna T, Hoff KJ, Lomsadze A, Stanke M, Borodovsky M. 2021. BRAKER2: Automatic eukaryotic genome annotation with GeneMark-EP+ and AUGUSTUS supported by a protein database. NAR Genom Bioinform 3:lqaa108.

Buchfink B, Xie C, Huson DH. 2015. Fast and sensitive protein alignment using DIAMOND. Nat Methods 12:59–60.

Bushmanova E, Antipov D, Lapidus A, Prjibelski AD. 2019. RnaSPAdes: a de novo transcriptome assembler and its application to RNA Seq data. GigaScience 8(9):giz100. 10.1093/gigascience/giz100

Cantalapiedra CP, et al. 2021. eggNOG-mapper v2: Functional annotation, orthology assignments, and domain prediction at the metagenomic scale. Mol Biol Evol 38:5825–5829.

Castelli M, Lanzoni O, Giovannini M, et al. 2022. ‘Candidatus Gromoviella agglomerans’, a novel intracellular Holosporaceae parasite of the ciliate Paramecium showing marked genome reduction. Environ Microbiol Rep 14:34–49.

Chang S, Sievert DM, Hageman JC, et al. 2003. Infection with vancomycin-resistant Staphylococcus aureus containing the vanA resistance gene. N Engl J Med 348:1342–1347.

Chen S, Zhou Y, Chen Y, Gu J. 2018. fastp: an ultra-fast all-in-one FASTQ preprocessor. Bioinformatics 34:i884–i890.

Clarke M, Lohan AJ, Liu B, et al. 2013. Genome of Acanthamoeba castellanii highlights extensive lateral gene transfer and early evolution of tyrosine kinase signaling. Genome Biol 14:R11. 10.1186/gb-2013-14-2-r11

Coale TH, Loconte V, Turk Kubo KA, et al. 2024. Nitrogen fixing organelle in a marine alga. Science 384(6692):217–222. 10.1126/science.adk1075

Chou S, Daugherty M, Peterson S, et al. 2015. Transferred interbacterial antagonism genes augment eukaryotic innate immune function. Nature 518:98–101.

Collingro A, Tischler P, Weinmaier T, et al. 2011. Unity in variety—the pan-genome of the Chlamydiae. Mol Biol Evol 28:3253–3270.

Conesa A, Goetz S, García-Gómez JM, Robles M, Talón M. 2004. Blast2GO: A universal annotation and visualization tool for functional genomics research. In: Spanish Bioinformatics Conference. p 211.

Criscuolo A, Gribaldo S. 2010. BMGE (Block Mapping and Gathering with Entropy): a new software for selection of phylogenetically informative regions from multiple sequence alignments. BMC Evol Biol 10:210.

Danecek P, Bonfield JK, Liddle J, et al. 2021. Twelve years of SAMtools and BCFtools. GigaScience 10(2):giab008. 10.1093/gigascience/giab008

Dirren S, Salcher MM, Blom JF, Schweikert M, Posch T. 2014. Ménage-à-trois: the amoeba Nuclearia sp. with its ecto- and endosymbiotic bacteria. Protist 165:745–758.

Dharamshi JE, Köstlbacher S, Schön ME, Collingro A, Ettema TJG, Horn M. 2023. Gene gain facilitated endosymbiotic evolution of Chlamydiae. Nat Microbiol 8:40–54. 10.1038/s41564-022-01284-9

Eddy SR. 2011. Accelerated profile HMM searches. PLoS Comput Biol 7:e1002195.

Edgcomb VP, Leadbetter ER, Bourland W, Beaudoin D, Bernhard JM. 2011. Structured multiple endosymbiosis of bacteria and archaea in a ciliate. Front Microbiol 2:55.

Edwards PA, Ericsson J. 1999. Sterols and isoprenoids: signaling molecules derived from the cholesterol biosynthetic pathway. Annu Rev Biochem 68:157–185. 10.1146/annurev.biochem.68.1.157

Eme L, Gentekaki E, Curtis B, Archibald JM, Roger AJ. 2017. Lateral gene transfer in the adaptation of the anaerobic parasite Blastocystis to the gut. Curr Biol 27:807–820.

Fehr A, et al. 2013. Candidatus Syngnamydia venezia, a novel member of the phylum Chlamydiae. PLoS ONE 8:e70853.

Filée J. 2014. Multiple occurrences of giant virus core genes acquired by eukaryotic genomes. Virology 466–467:1–9.

Floriano AM, Castelli M, Krenek S, et al. 2018. Genome sequence of Candidatus Fokinia solitaria. Genome Biol Evol 10:1120–1126.

Garushyants SK, Beliavskaia AY, Malko DB, et al. 2018. Comparative genomic analysis of Holospora spp. Front Microbiol 9:738.

George EE, et al. 2020. Highly reduced genomes of protist endosymbionts show evolutionary convergence. Curr Biol 30:925–933.e3.

Gouy M, Tannier E, Comte N, Parsons DP. 2021. SeaView version 5: A multiplatform software for multiple sequence alignment, molecular phylogenetic analyses, and tree reconciliation. In: Katoh K, editor. Methods Mol Biol, vol 2231. Humana Press. p 241–260. 10.1007/978-1-0716-1036-7_15

Gurevich A, Saveliev V, Vyahhi N, Tesler G. 2013. QUAST: quality assessment tool for genome assemblies. Bioinformatics 29:1072–1075.

Gutiérrez G, Chistyakova LV, Villalobo E, Kostygov AY, Frolov AO. 2017. Identification of Pelomyxa palustris endosymbionts. Protist 168(4):408–424. 10.1016/j.protis.2017.06.001

Haas BJ, Papanicolaou A. 2022. TransDecoder (Find Coding Regions Within Transcripts). GitHub. https://github.com/TransDecoder/TransDecoder

Hamann E, Gruber-Vodicka H, Kleiner M, et al. 2016. Environmental Breviatea harbour mutualistic Arcobacter epibionts. Nature 534:254–258.

Hehenberger E, Burki F, Kolisko M, Keeling PJ. 2016. Functional relationship between a dinoflagellate host and its diatom endosymbiont. Mol Biol Evol 33:2376–2390.

Henze K, Martin W, Schnarrenberger C. 2002. Endosymbiotic gene transfer. In: Horizontal Gene Transfer, 2nd ed. Academic Press.

Hoff KJ, Lomsadze A, Borodovsky M, Stanke M. 2019. Whole-genome annotation with BRAKER. Methods Mol Biol 1962:65–95.

Hoang DT, Chernomor O, von Haeseler A, Minh BQ, Vinh LS. 2018. UFBoot2. Mol Biol Evol 35:518–522.

Hugoson E, Guliaev A, Ammunét T, Guy L. 2022. Host adaptation in Legionellales is 1.9 Ga, coincident with eukaryogenesis. Mol Biol Evol 39(3):msac037. 10.1093/molbev/msac037

Husnik F, McCutcheon JP. 2018. Functional horizontal gene transfer from bacteria to eukaryotes. Nat Rev Microbiol 16:67–79.

Husník F, McCutcheon JP. 2017. Functional horizontal gene transfer from bacteria to eukaryotes. Nat Rev Microbiol 16(2):67–79. 10.1038/nrmicro.2017.137

Husnik F, Tashyreva D, Boscaro V, George EE, Lukeš J, Keeling PJ. 2021. Bacterial and archaeal symbioses with protists. Curr Biol 31(13):R862–R877. 10.1016/j.cub.2021.05.049

Ishida K, Sekizuka T, Hayashida K, et al. 2014. Amoebal endosymbiont Neochlamydia genome sequence illuminates the bacterial role in the defense of the host amoebae against Legionella pneumophila. PLoS ONE 9(4):e95166. 10.1371/journal.pone.0095166

Jones P, Binns D, Chang HY, et al. 2014. InterProScan 5. Bioinformatics 30:1236–1240.

Katoh K, Standley DM. 2013. MAFFT version 7. Mol Biol Evol 30:772–780.

Keeling PJ. 2024. Horizontal gene transfer in eukaryotes. Nat Rev Genet.

Keeling PJ, McCutcheon JP. 2017. Endosymbiosis: the feeling is not mutual. J Theor Biol 434:75–79.

Keeling PJ, McCutcheon JP, Doolittle WF. 2015. Symbiosis becoming permanent: Survival of the luckiest. Proc Natl Acad Sci USA 112(33):10101–10103. 10.1073/pnas.1513346112

Keeling PJ, Palmer JD. 2008. Horizontal gene transfer in eukaryotic evolution. Nat Rev Genet 9:605–618.

Khalil JYB, Benamar S, Baudoin JP, et al. 2016. Developmental cycle and genome analysis of Rubidus massiliensis, a new Vermamoeba vermiformis pathogen. Front Cell Infect Microbiol 6:31. 10.3389/fcimb.2016.00031

Koraimann G, Wagner MA. 2014. Social behavior and decision making in bacterial conjugation. Front Cell Infect Microbiol 4:54. 10.3389/fcimb.2014.00054

Kostygov AY, Butenko A, Nenarokova A, et al. 2017. Identification of Pelomyxa palustris endosymbionts. Protist 168(4):408–424. 10.1016/j.protis.2017.06.001

Larkum AWD, Lockhart PJ, Howe CJ. 2007. Shopping for plastids. Trends Plant Sci 12:189–195.

Latorre A, Manzano Marín A. 2017. Dissecting genome reduction and trait loss in insect endosymbionts. Ann NY Acad Sci 1389(1):52–75. 10.1111/nyas.13222

Leander BS, Keeling PJ. 2004. Symbiotic innovation in the oxymonad Streblomastix strix. J Eukaryot Microbiol 51(3):291–300. 10.1111/j.1550-7408.2004.tb00569.x

Lee MC, Marx CJ. 2012. Repeated genome reduction of accessory genes. PLoS Genet 8:e1002651.

Li H, Durbin R. 2009. Fast and accurate short read alignment with Burrows-Wheeler transform. Bioinformatics 25:1754–1760.

Liechti N, Schürch N, Bruggmann R, Wittwer M. 2019. Nanopore sequencing improves the draft genome of the human pathogenic amoeba Naegleria fowleri. Sci Rep 9:16040. 10.1038/s41598-019-52572-0

Maki T, Yoshinaga I, Katanozaka N, Imai I. 2004. Phylogenetic analysis of intracellular bacteria. Aquat Microb Ecol 36:123–135.

Manni M, Berkeley MR, Seppey M, Simão FA, Zdobnov EM. 2021. BUSCO update. Mol Biol Evol 38:4647–4654.

McCutcheon JP, Moran NA. 2011. Extreme genome reduction in symbiotic bacteria. Nat Rev Microbiol 10:13–26.

Minh BQ, Schmidt HA, Chernomor O, et al. 2020. IQ-TREE 2: New models and efficient methods for phylogenetic inference in the genomic era. Mol Biol Evol 37(5):1530–1534. 10.1093/molbev/msaa015

Mira A, Ochman H, Moran NA. 2001. Deletional bias and the evolution of bacterial genomes. Trends Genet 17:589–596.

Moran NA. 2002. Microbial minimalism. Cell 108:583–586.

Moran Y, Fredman D, Szczesny P, Grynberg M, Technau U. 2012. Recurrent horizontal transfer of bacterial toxin genes to eukaryotes. Mol Biol Evol 29:2223–2230.

Nawrocki EP, Eddy SR. 2013. Infernal 1.1: 100 fold faster RNA homology searches. Bioinformatics 29(22):2933–2935. 10.1093/bioinformatics/btt509

Nikoh N, et al. 2010. Bacterial genes in the aphid genome. PLoS Genet 6:e1000827.

Nowack ECM. 2014. Paulinella chromatophora — rethinking the transition from endosymbiont to organelle. Acta Soc Bot Pol 83(4):387–397. 10.5586/asbp.2014.049

Nugraha RYB, Jeelani G, Nozaki T. 2022. GABA metabolism in parasitic protozoa. Trends Parasitol 38:462–477.

Nylund A, Pistone D, Trösse C, et al. 2018. Genotyping of Candidatus Syngnamydia salmonis (Chlamydiales; Simkaniaceae) co cultured in Paramoeba perurans (Amoebozoa; Paramoebidae). Arch Microbiol 200(6):859–867. 10.1007/s00203-018-1488-0

Pagnier I, Yutin N, Croce O, et al. 2015. Babela massiliensis, a representative of a widespread bacterial phylum with unusual adaptations to parasitism in amoebae. Biol Direct 10:13. 10.1186/s13062-015-0043-z

Patro R, Duggal G, Love MI, Irizarry RA, Kingsford C. 2017. Salmon. Nat Methods.

Price MN, Dehal PS, Arkin AP. 2009. FastTree. Mol Biol Evol 26:1641–1650.

Prieto Santos MI, Martin Checa J, Balaña Fouce R, Garrido Pertierra A. 1986. A pathway for putrescine catabolism in Escherichia coli. Biochim Biophys Acta 880(2–3):242–244. 10.1016/0304-4165(86)90085-1

Prokopchuk G, Tashyreva D, Yabuki A, et al. 2019. Morphological, ultrastructural, motility and evolutionary characterization of two new Hemistasiidae species. Protist 170(3):259–282. 10.1016/j.protis.2019.04.001

Qin QL, et al. 2019. Genomic reduction in Idiomarina. mBio 10:e02545–18.

Read TD, et al. 2013. Chlamydia psittaci comparative genomics. mBio 4:e00604–12.

Romero M, Cerritos R, Ximenez C. 2016. Horizontal gene transfers from bacteria to Entamoeba complex: a strategy for dating events along species divergence. J Parasitol Res 2016:3241027. 10.1155/2016/3241027

Schaack S, Gilbert C, Feschotte C. 2010. Promiscuous DNA. Trends Ecol Evol 25:537–546.

Shiohama Y, et al. 2022. Candidatus Hydrogeosomobacter endosymbioticus. Microbiol Resour Announc 11:e01150.

Simão FA, et al. 2015. BUSCO. Bioinformatics 31:3210–3212.

Song W, Wemheuer B, Zhang S, et al. 2019. MetaCHIP. Microbiome 7:36.

Takeuchi M, Kuwahara H, Murakami T, et al. 2020. Parallel reductive genome evolution in Desulfovibrio ectosymbionts independently acquired by Trichonympha protists in the termite gut. ISME J 14(9):2288–2301. 10.1038/s41396-020-0688-1

Tashyreva D, Prokopchuk G, Votýpka J, et al. 2018. Life cycle, ultrastructure, and phylogeny of new diplonemids and their endosymbiotic bacteria. mBio 9(2):e02447–17. 10.1128/mBio.02447-17

Tashyreva D, Simpson AGB, Prokopchuk G, et al. 2022. Diplonemids – A review on “new” flagellates on the oceanic block. Protist 173(2):125868. 10.1016/j.protis.2022.125868

Tashyreva D, Votýpka J, Yabuki A, Horák A, Lukeš J. 2025. Description of new diplonemids (Diplonemea, Euglenozoa) and their endosymbionts: charting the morphological diversity of these poorly known heterotrophic flagellates. Protist 177:126090. 10.1016/j.protis.2025.126090

Thomas V, Casson N, Greub G. 2006. Criblamydia sequanensis. Environ Microbiol 8:2125–2135.

Trapnell C, Pachter L, Salzberg SL. 2009. TopHat: discovering splice junctions with RNA Seq. Bioinformatics 25(9):1105–1111. 10.1093/bioinformatics/btp120

Valach M, Moreira S, Petitjean C, et al. 2023. Recent expansion of metabolic versatility in Diplonema papillatum, the model species of a highly speciose group of marine eukaryotes. BMC Biol 21(1):99. 10.1186/s12915-023-01563-9

Van Etten J, Bhattacharya D. 2020. Horizontal gene transfer in eukaryotes. Trends Genet 36:915–925.

Vannini C, Rosati G, Verni F, Petroni G. 2004. Candidatus Devosia euplotis. Int J Syst Evol Microbiol 54:1151–1156.

Verster KI, Tarnopol RL, Akalu SM, Whiteman NK. 2021. Horizontal transfer of microbial toxin genes to gall midge genomes. bioRxiv.

von der Heyden S, Chao EE, Vickerman K, Cavalier-Smith T. 2004. Ribosomal RNA phylogeny of bodonid and diplonemid flagellates and the evolution of Euglenozoa. J Eukaryot Microbiol 51(4):402–416. 10.1111/j.1550-7408.2004.tb00387.x

Wang HC, Minh BQ, Susko E, Roger AJ. 2018. Modeling site heterogeneity. Syst Biol 67:216–235.

Wang S, Luo H. 2021. Dating Alphaproteobacteria evolution with eukaryotic fossils. Nat Commun 12:3324. 10.1038/s41467-021-23645-4

Zhan SH, French L. 2019. Morphine biosynthesis enzyme similarity searches. J Med Microbiol 68:952–956.

